# BRCA2 chaperones RAD51 to single molecules of RPA-coated ssDNA

**DOI:** 10.1101/2022.12.28.522061

**Authors:** Jason C. Bell, Christopher C. Dombrowski, Jody L. Plank, Ryan B. Jensen, Stephen C. Kowalczykowski

## Abstract

Mutations in the breast cancer susceptibility gene, BRCA2, greatly increase an individual’s lifetime risk of developing breast and ovarian cancers. BRCA2 suppresses tumor formation by potentiating DNA repair via homologous recombination. Central to recombination is the assembly of a RAD51 nucleoprotein filament, which forms on single-stranded DNA (ssDNA) generated at or near the site of chromosomal damage. However, Replication Protein-A (RPA) rapidly binds to and continuously sequesters this ssDNA, imposing a kinetic barrier to RAD51 filament assembly that suppresses unregulated recombination. Recombination mediator proteins––of which BRCA2 is the defining member in humans ––alleviate this kinetic barrier to catalyze RAD51 filament formation. We combined microfluidics, microscopy, and micromanipulation to directly measure both the binding of full-length BRCA2 to––and the assembly of RAD51 filaments on––a region of RPA-coated ssDNA within individual DNA molecules designed to mimic a resected DNA lesion common in replication-coupled recombinational repair. We demonstrate that a dimer of RAD51 is minimally required for spontaneous nucleation; however, growth self-terminates below the diffraction limit. BRCA2 accelerates nucleation of RAD51 to a rate that approaches the rapid association of RAD51 to naked ssDNA, thereby overcoming the kinetic block imposed by RPA. Furthermore, BRCA2 eliminates the need for the rate-limiting nucleation of RAD51 by chaperoning a short pre-assembled RAD51 filament onto the ssDNA complexed with RPA. Therefore, BRCA2 regulates recombination by initiating RAD51 filament formation.

**Significance:** Despite decades of genetic and cell biological studies, mechanistic biochemical analyses of human BRCA2 function in recombinational DNA repair have only been possible since the purification of full-length BRCA2. These mechanistic studies crucially inform with respect to the molecular function of BRCA2 in genome maintenance. Here, we use single-molecule methods to visualize the assembly of RAD51 on individual RPA-coated ssDNA molecules and to see how this process is regulated by the tumor suppressor protein, BRCA2. We show that BRCA2 serves as a chaperone to nucleate RAD51 and deliver it to RPA-coated ssDNA. This work advances understanding of the molecular functions of BRCA2 and, consequently, the molecular etiology of breast cancer in an important way.

## Introduction

Mutations in the breast cancer susceptibility gene, *BRCA2*, greatly increase the lifetime risk of developing breast and ovarian cancers (1). *BRCA2* was identified in families with a high incidence of breast cancer (2). The relationship between BRCA2 and homologous recombination became evident when its interaction with RAD51 was discovered (3, 4). Cells lacking BRCA2 function suffer from genomic instability, loss of DNA damage-induced RAD51 foci, and extreme sensitivity to crosslinking agents such as mitomycin-C (MMC) and cisplatin (5-8). BRCA2 suppresses tumor formation by potentiating DNA repair via homologous recombination (6, 9-12) (13).

As mentioned above, central to recombination is assembly of the RAD51 nucleoprotein filament onto ssDNA generated at the site of chromosomal damage. However, RPA rapidly binds to ssDNA, kinetically impeding RAD51 filament assembly. BRCA2, which is the defining member of recombination mediators in humans (9, 11, 14), alleviates this kinetic barrier to catalyze RAD51 filament formation (15, 16). Structurally, a unique feature of human BRCA2 is the presence of eight conserved BRC repeat sequences (3, 17, 18). Each BRC repeat binds RAD51 *via* an interface that mimics the RAD51 interface (3, 19-21). The carboxy-terminus of BRCA2 possesses three OB (oligonucleotide binding) folds with a tower structure protruding out of the second OB fold (22). This juxtaposition of DNA binding regions and the internal repeats necessary for RAD51 binding, suggested a model whereby BRCA2 binds ssDNA or an ssDNA/dsDNA junction and loads RAD51 onto the ssDNA (23).

*In vitro*, full-length human BRCA2 promotes assembly of RAD51 onto ssDNA complexed with RPA, making it a *bona fide* mediator (9, 11, 24). BRCA2 binds to ssDNA with high affinity (∼nM) in a structure-independent manner (i.e., an ssDNA-dsDNA junction is not required). BRCA2 also binds at least 6 monomers of RAD51, and likely up to 8, in a species-specific manner via its 8 BRC repeats (9). The BRC repeats are neither identical in sequence nor in function. BRCA2 binds RAD51 through its eight BRC repeats in two distinct ways. BRC repeats 1-4 bind to free RAD51 with a high affinity and block both ssDNA-dependent ATP hydrolysis and dsDNA binding (9, 20, 21). In contrast, BRC repeats 5-8 bind to and stabilize the nascent RAD51-ssDNA filament to prevent disassembly (21, 25). BRC1-4 also block ATP hydrolysis by RAD51, which prevents dissociation of RAD51 from ssDNA and keeps RAD51 in the active ATP-bound form. This partitioning of labor suggested a mechanism where BRCA2 delivers up to 4 molecules of RAD51 to the ssDNA to serve as the nucleus to initiate assembly, and then the next four BRC repeats stabilize the next 4 molecules of RAD51 as they bind to ssDNA adding to the nucleus. In this way, BRCA2 was hypothesized to chaperone nascent filament assembly of 8 RAD51 monomers, which comprises slightly more than one turn of the filament (∼6 monomers).

In this work, we used optical trapping to physically manipulate single molecules of DNA that were engineered to contain a large ssDNA gap, mimicking a replication gap that is one of several physiological substrates for the recombinational DNA repair. We incubated these gapped molecules with RPA and then, using optical trapping, we “dipped” each molecule into a solution containing RAD51 with, or without, purified BRCA2. By measuring the binding locations of each RAD51 and BRCA2, and the concentration-dependence of the kinetics of RAD51 nucleation on RPA-coated ssDNA, we ascertained that BRCA2 functions as a molecular chaperone for RAD51 to accelerate filament nucleation to overcome the kinetic inhibition imposed by RPA.

## Results and Discussion

Using a microfluidic multi-channel flow cell, coupled with a fluorescence microscope and a dual optical trap, two streptavidin-coated beads were isolated and a single molecule of DNA, biotinylated at each end, was tethered between the beads *in situ* (Fig. 1A, *illustration*) (15, 26-28). To image the assembly of RAD51 filaments on RPA-coated ssDNA, we used a dsDNA substrate containing an ssDNA gap of 8,155 nucleotides (nt) flanked by 21,080 bp and 24,590 bp of dsDNA (Fig. 1B) (28). To create this substrate, we used a derivative of bacteriophage λgt11 containing a λC31 *attP* recognition site and a derivative of bacteriophage M13mp7 ssDNA containing the *attB* sequence, which creates the duplex DNA recognition site upon annealing of a complementary DNA strand. Integration of the circular ssDNA into biotinylated λ DNA resulted in a DNA substrate that we hereafter refer to as gapped λ DNA. This substrate mimics a replication gap––one of the primary lesions requiring recombination-coupled DNA repair in normal cell division––arising from incomplete lagging strand synthesis or resection from a replication block. Unless specifically indicated, the gapped λ DNA was pre-incubated with purified human RPA before capture in the optical traps, and RPA was present in each channel of the flow cell to ensure that ssDNA within the gap would be entirely coated with RPA.

**Figure 1:**
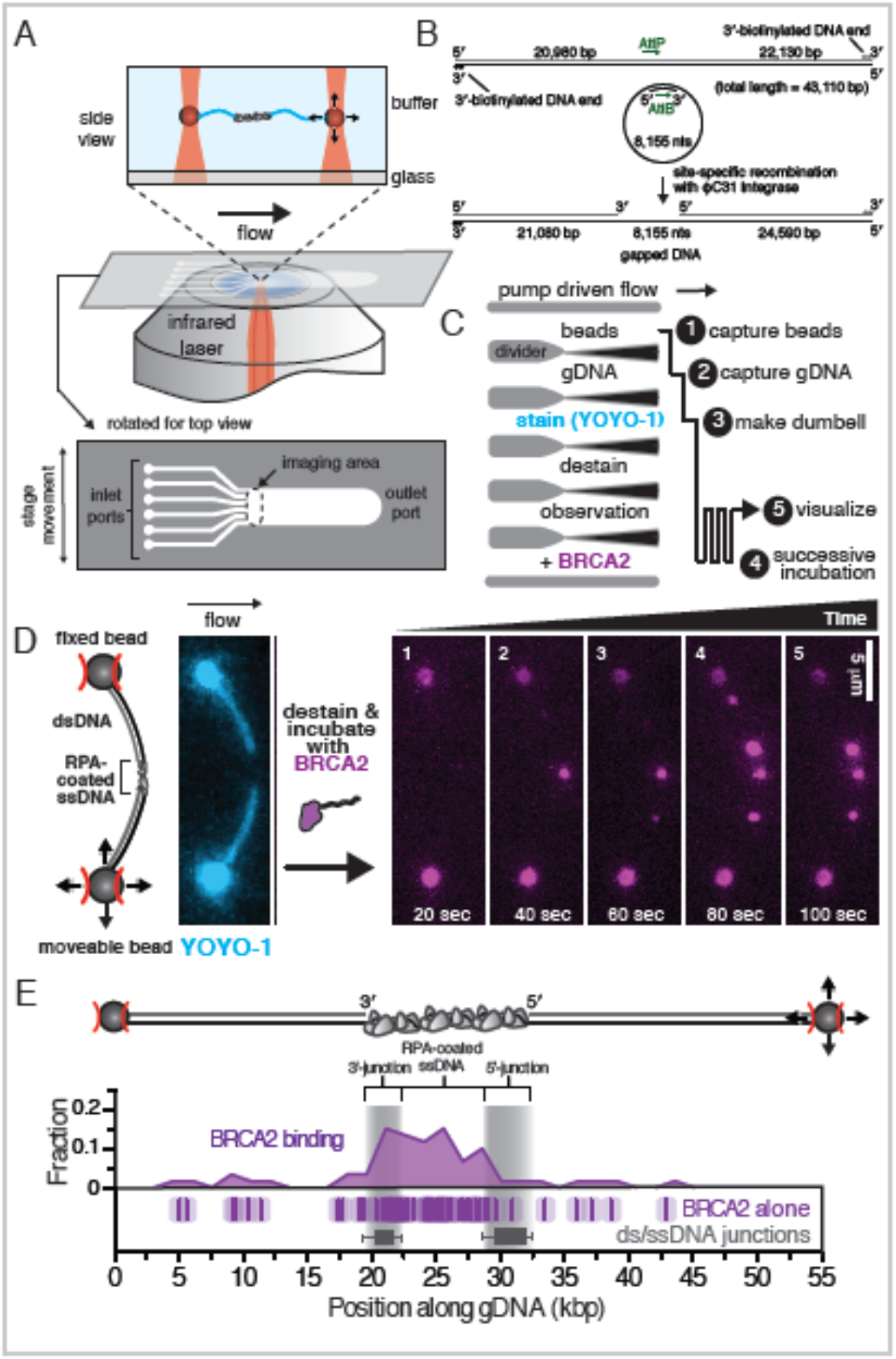
Direct imaging of BRCA2 binding to RPA-coated ssDNA on single molecules of gapped λ DNA. **A)** Schematic of experimental approach combining fluorescence microscopy, a microfluidic flow-cell, and optical-trapping, as well as the micromanipulation used to capture and image BRCA2 on individual DNA molecules. **B)** Illustration of the gapped λ DNA generated through *in vitro* recombination of circular ssDNA with an engineered λ DNA. **C)** Schematic of experimental protocol: each molecule of gapped λ DNA was captured and micromanipulated between two beads held in separately controllable optical traps. The molecule was moved between solutions in a six-channel flow cell, and successively incubated in a solution containing BRCA2. **D)** Cartoon and microscopic image of a single-molecule of gapped λ DNA (*left, stained with YOYO-1, cyan*) that was destained and then successively incubated with BRCA2 (5 nM) plus α-BRCA2 and α-IgG^AF546^. Montage shows BRCA2 (*magenta*) binding to the gapped λ DNA at increasing time intervals. **E)** Cartoon representation of the gapped λ DNA between two beads (*top*) and histogram (*middle*) of binding positions of BRCA2 (number of foci, *N*=60). Each data point is also plotted as a single tick (*bottom*) where the semi-transparent box represents the standard error associated with assigning position owing to the optical resolution of the microscope. Gray bars represent the 10-90^th^ percentile range of the 5’- and 3’-termined junctions (*N*=98).

Each isolated molecule was first visualized using YOYO-1 to stain the dsDNA portions, so as to establish the integrity and orientation of each molecule. YOYO-1 was then dissociated by incubation in the “destain” channel, and each molecule was then successively incubated in a channel containing full-length human BRCA2 (purified as previously described (9)), and visualized by co-incubation with a fluorescently labeled antibody (Fig. 1C). Each molecule was rotated perpendicular to the flow to maximize spatial resolution, while minimizing flow-exerted forces on the RPA-coated ssDNA. With time, BRCA2 binding was directly observed along the DNA (Fig. 1D and Movie 1). The initial position of each binding event was determined by measuring the relative distance of each fluorescent focus along the contour length of the DNA between the beads. Due to intercalation of YOYO-1 into dsDNA, the contour length of the molecule changes after YOYO-1 dissociation but we could determine the orientation of each molecule and the position of each binding event relative to the ssDNA-dsDNA junctions, which is plotted in Fig. 1E. In aggregate, BRCA2 binding demonstrated a preference for binding the ssDNA region, despite the presence of RPA, consistent with our previous observation that BRCA2 preferentially binds ssDNA over dsDNA^2^. Binding events observed on dsDNA were more labile, and infrequently, molecules of BRCA2 bound to dsDNA under flow were observed sliding on the dsDNA (Movie 1 and Supplementary Figure 1).

To visualize formation of a RAD51 filament, we attached gapped λ DNA to the surface of a PEG-coated flow cell and initially used total internal reflection fluorescence (TIRF) microscopy (Fig. 2B), taking advantage of the higher throughput of TIRF experiments (28). The dsDNA region of the molecule was imaged using SYTOX Orange. The ssDNA gap was clearly visible owing to the absence of SYTOX Orange staining. When purified fluorescent RAD51 (previously described) (29) was injected into the flow cell, the RAD51-binding buffer caused dissociation of the SYTOX Orange from the dsDNA within 1-3 seconds (s) and, with time, RAD51 filaments formed on the ssDNA region. In the absence of RPA, association of RAD51 was almost as fast as we could measure: the lag time for nucleation––determined as the time for detection of the first diffraction-limited focus on the ssDNA––was on the order of 5-10 s, and binding was complete after 1-2 minutes (Fig. 2B and Movie 2). When RPA was included, RAD51 filament formation was potently blocked (Fig. 2C): in this example, nucleation was not evident until 120 s and complete filaments (ascertained by ‘filling in’ of the gap) were not observed even at 240 s (Fig. 2c and Movie 3). It is worth noting that previous single-molecule experiments demonstrated the binding of RAD51 to dsDNA in the absence of salt; however, this off-target binding is dramatically reduced between 150-300 mM NaCl (29), the range of ionic strength physiologically relevant for the mammalian nucleus (30). Therefore, all RAD51 binding experiments were performed in 200 mM NaCl (*see Methods*).

**Figure 2:**
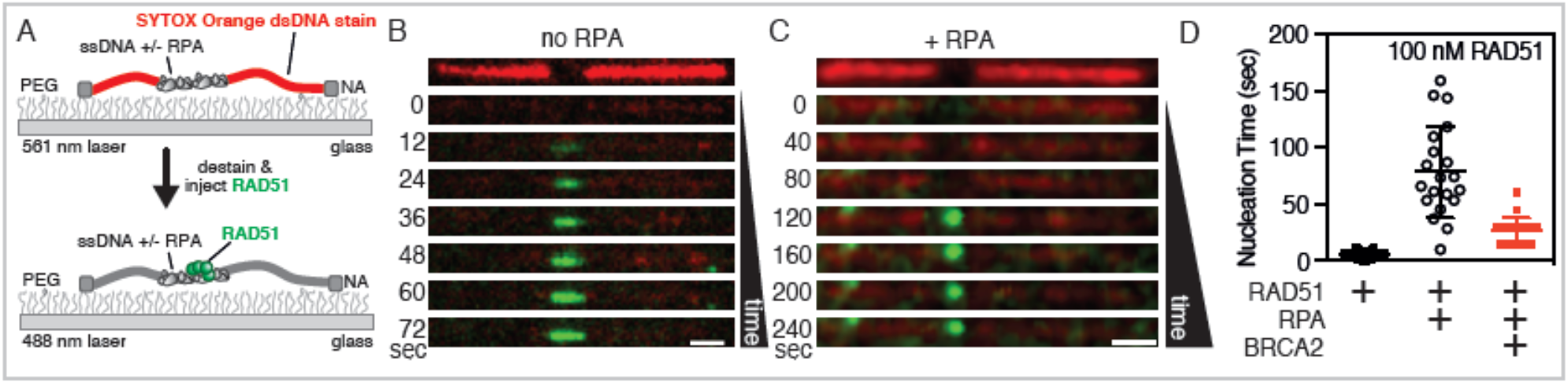
Direct imaging of RAD51 nucleation and filament formation in the absence and presence of RPA. **A)** Schematic of a single molecule of gapped λ DNA attached at each end to a PEG-coated surface via neutravidin. The dsDNA region was initially visualized using SYTOX Orange (*red*), which was subsequently dissociated upon the addition of binding buffer and fluorescein-RAD51 (*green*). **B)** In the absence of RPA, fluorescent RAD51 rapidly filled the ssDNA region. **C)** When RPA was present, RAD51 binding was slower and punctate. **D)** Comparison of the lag time in the absence (black filled symbols) or presence (black open symbols) of RPA measured using TIRF microscopy. Lag times in the presence of BRCA2 were measured using optical trapping (*see text*) and are shown in red symbols for comparison. Lines represent the arithmetic mean and error bars represent standard deviation. No RPA: 5±3 (*N*=111); +RPA: 78±40 (*N*=20); +RPA/BRCA2: 27±11 (*N*=22). Scale bar in panels *b* and *c* is 2 µm.

We next measured the times for RAD51 nucleation in the absence and presence of RPA. On average, nucleation of RAD51 in the absence of RPA was fast, limited primarily by the dead-time of the injection of fluorescent RAD51 into our flow cell, with an average lag time of 5±3 s (Fig. 2D, *leftmost*). In the presence of RPA, the nucleation time slowed by 16-fold, with an average lag time of 78±46 s (Fig. 2D, *center*). The lag times obtained from our TIRF and trapping assays were identical, within error (Supplementary Figure 2); however, we elected to use the optical trapping assay for further experiments containing BRCA2 owing to the smaller volumes and lower flow rates, which greatly reduced the material required for each experiment. When we measured the nucleation time for RAD51 in the presence of BRCA2, we observed a 3-to 4-fold net reduction from 78±40 s in the absence of BRCA2, to 27±11 s in the presence of BRCA2 (Fig. 2D, *rightmost*) which approaches, within a factor of four, the nucleation rate observed when RPA is absent.

We subsequently visualized RAD51 nucleation on gapped λ DNA using two-color fluorescence microscopy (Fig. 3A, *schematic*), to simultaneously image both RAD51 nucleation and BRCA2 binding (Fig. 3A). These experiments confirmed the expected co-localization of RAD51 and BRCA2 in 56% of these RAD51 nucleation events, slightly more than the 40% expected due to the ∼2.5-fold excess of RAD51 over binding sites on BRCA2. As previously described for Figure 1, we measured the position of each nascent RAD51 filament relative to the junctions along the gapped λ DNA and plotted every position observed for the duration of the imaging protocol (Fig. 3C), in either the absence (*green)* or presence (*red)* of BRCA2 in the absence of antibody so as not to potentially alter the outcome. RAD51 nucleation was exclusively observed within the RPA-coated ssDNA region, with usually one RAD51 focus per DNA molecule and generally no more than three. There was an apparent skewed preference (Fig. 3D) for the ssDNA region near the 3’-terminated junction relative to the 5’-junction (*odds ratio*=4.7, *p*=0.3377; Fisher’s exact test). In the presence of BRCA2, we observed a 6.3-fold increase in the number of nucleation events near the 5’-terminated junction relative to the RAD51-alone condition (*odds ratio*=6.3, *p*=0.1113; Fisher’s exact test) resulting in nascent RAD51 foci forming closer to the edges of the RPA-coated ssDNA region, consistent with both the 2-fold higher affinity for ssDNA-dsDNA junctions established from ensemble studies (9) and a potential kinetic bias resulting from BRCA2 sliding down the dsDNA under flow.

**Figure 3:**
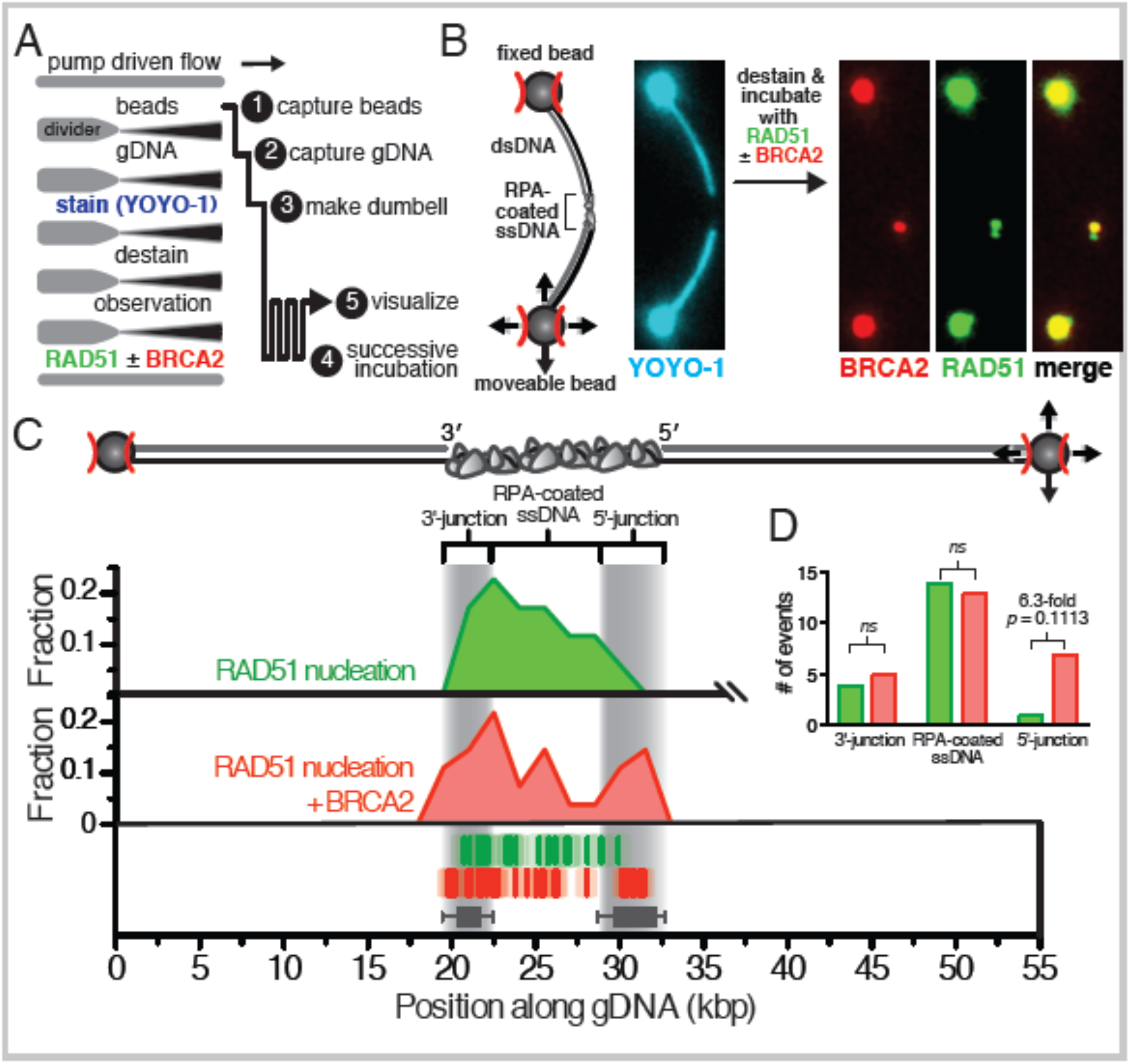
RAD51-BRCA2 complexes, in contrast to BRCA2 alone, are focused to the ssDNA regions. **A)** Schematic of optical-trap experiments designed to visualize nucleation of RAD51 on gapped λ DNA in the absence or presence of BRCA2. **B)** A single molecule of gapped λ DNA with bound RPA held between two optical traps and co-incubated with 100 nM RAD51 (green) and 5 nM BRCA2 (red, α-MBP^AF546^). **C)** Cartoon of the gapped λ DNA (*top*) and histogram (*middle*) showing positions of all RAD51 nucleation events either alone (*green*) (*N*=18) or when co-incubated with BRCA2 (*red*) in the absence of antibody (*N*=28). Positions of individual foci are plotted (*bottom*) relative to position of the 5’- and 3’-termined junctions. The transparent box around each dash represents the standard error owing to the optical resolution of our microscope. Gray bars represent the 10-90^th^ percentile range of the 5’- and 3’-termined junctions (*N*=98). **D)** Bar plot of the number of RAD51 nucleation events in the absence (*green*) or presence of BRCA2 (*red*) observed in the regions nearest the 3’-junction, clearly in the middle of the RPA-coated region, or near the 5’-junction. The odds ratios and p-values were calculated using Fisher’s exact test.

We next analyzed the kinetics of RAD51 nucleation on RPA-coated ssDNA as a function of RAD51 concentration, plotting the cumulative frequency of RAD51 nucleation events as a function of increasing incubation time (Fig. 4A and 4B). When analyzed in this way, the observed cumulative frequency increases exponentially with time, and a characteristic nucleation time can be determined by fitting the data to a single exponential curve. As expected, the nucleation time decreased as the concentration of RAD51 increased. In the absence of BRCA2 (Fig. 4A), spontaneous nucleation is strongly dependent on the concentration of RAD51, where the rate of nucleation is proportional to the concentration of the filament forming protein raised to the power of the minimum nucleation species (i.e., *J* µ *k*•[RAD51]^*n*^, where *J* = 1/lag time, *n* is the size of the oligomer and *k* is a dimensionless rate constant) (28, 29, 31, 32). Using this kinetic analysis, we observed a power dependence where *n* = 1.5±0.3 (Fig. 4C), indicating that RAD51 nucleation proceeds through either monomers or dimers; however, since RAD51 nucleation is nucleotide dependent and the nucleotide binding interface lies between two half sites formed between adjacent monomers, we conclude that in the absence of BRCA2, the smallest oligomer required for a single nucleation event on RPA-coated ssDNA is a dimer of RAD51, similar to previous observations with both RecA (28) and RAD51 (29, 32).

**Figure 4:**
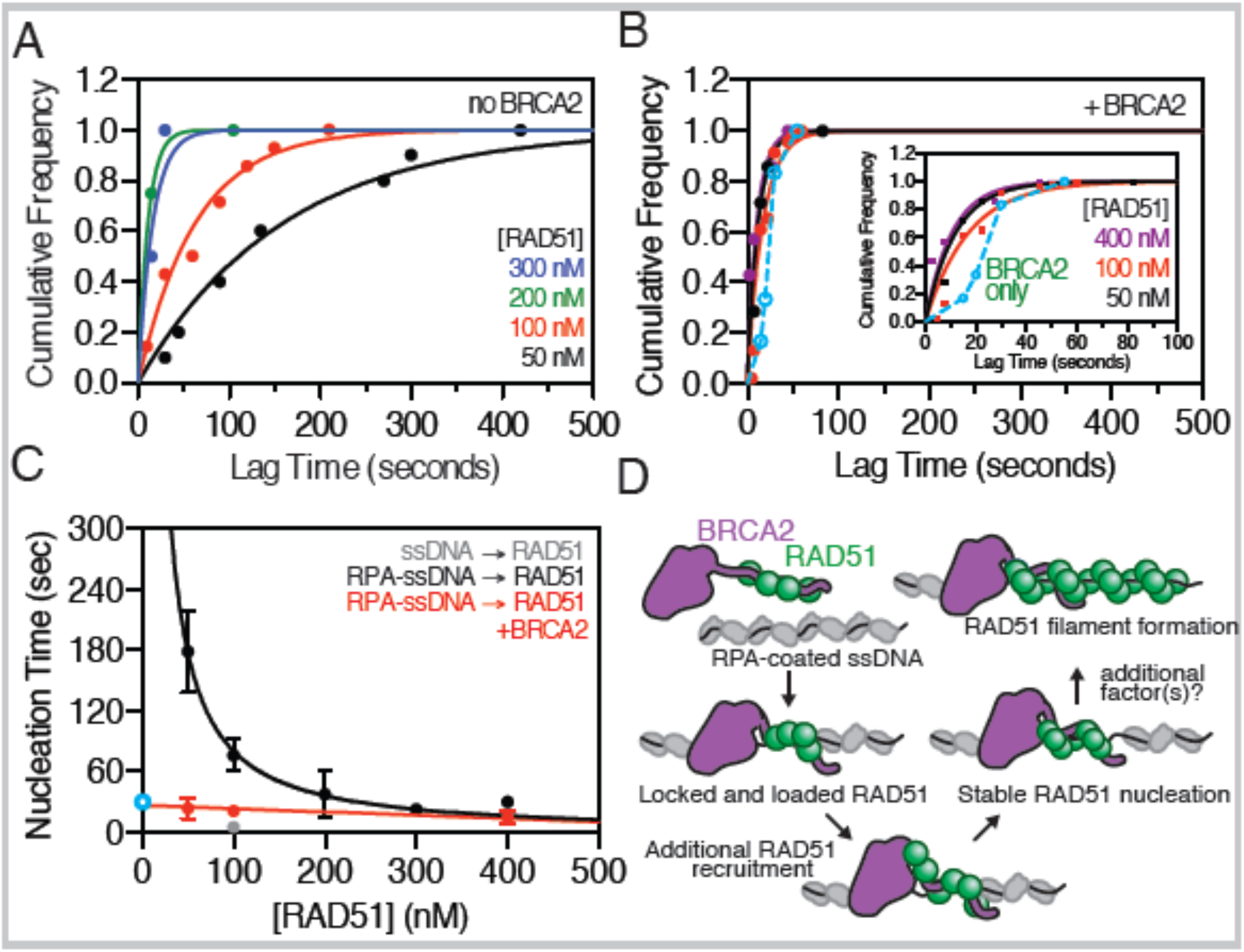
Nucleation of RAD51 requires a dimer and is accelerated by BRCA2 by delivering a pre-nucleated complex. **A)** The cumulative frequency of RAD51 nucleation measured by optical trapping is plotted as a function of increasing incubation time at varying concentrations of RAD51 in the absence or **B)** presence of BRCA2. Inset shows a zoomed in scale. The solid lines represent non-linear fitting to a single-exponential rate equation: 50 nM: t_1/2_=110±10 s, 100 nM: t_1/2_=47±4 s, 200 nM: t_1/2_=8±1 s, 300 nM: t_1/2_=11±4 s (std. error). The open symbols and cyan dashed line represent the kinetics of BRCA2 binding in the absence of RAD51 as measured in Figure 1 and is shown as a comparison to the kinetics of RAD51 nucleation (*see text*). **C)** The arithmetic mean nucleation time plotted as a function of increasing RAD51 concentration in the absence: 50 nM: 179±41 s (*N*=10), 100 nM: 76±16 s (*N*=14), 200 nM: 38±23 s (*N*=4), 300 nM: 23±4 s (*N*=4), 400 nM: 30±6 s (*N*=6), or presence of BRCA2: 50 nM: 24±10 s (*N*=7), 100 nM: 21±2 s (*N*=46), 400 nM: 15±6 s (*N*=7). The black curve represents a fit to the power law (***J***=***k***[RAD51]^***n***^) where ***n*** = 1.5±0.3 (std. error). The cyan symbol represents the half-time for BRCA2 binding in the absence of RAD51, measured with α-BRCA2 plus α-IgG^AF546^: 30±6 s (*N*=6). The red line is for visual purposes only; gray symbol is the rate in the absence of RPA: 5.2±0.3 s (*N*=111). Error and error bars are standard error of the mean, and if not visible, are smaller than the symbol. **D)** Model of BRCA2-mediated RAD51 nucleation on RPA-coated ssDNA.

When we repeated the experiment in the presence of BRCA2, the RAD51 nucleation times were decreased (Fig. 4B), as anticipated. Unexpectedly, the kinetics of RAD51 nucleation were independent of RAD51 concentration across the accessible range of our assay (Fig. 4C). To determine whether the time to RAD51 nucleation was strictly limited by the kinetics of BRCA2 binding to the ssDNA coated with RPA, we measured the kinetics of BRCA2 binding to RPA-coated ssDNA in the absence of RAD51 as shown in Fig. 1D, and plotted the data (Fig. 4B, *blue open circles & dashed line, BRCA2 only*). The binding of BRCA2 in the absence of RAD51 exhibits a sigmoidal increase with a characteristic half-time of ∼25 s. In contrast, nucleation of RAD51 in the presence of BRCA2 was ∼two-fold faster than binding of BRCA2 alone (Fig. 4C, *purple*), with a characteristic half-time of 7-12 s and was independent of the concentration of RAD51 (Fig. 4C, *red*). The faster rate of DNA binding for BRCA2 in the presence of RAD51 indicates that the proteins bind as a complex, consistent with the role of BRCA2 functioning as a molecular chaperone to promote RAD51 nucleation. The conclusion is quantitatively substantiated by the data in Fig 4C which reveal that BRCA2 eliminates the apparent concentration dependence of RAD51 nucleation, demonstrating that RAD51 is delivered by BRCA2 to the ssDNA in a pre-nucleated form and bypassing the otherwise rate-limiting step of dimer-formation during filament assembly.

Our observations advance a model whereby BRCA2 functions to chaperone RAD51 to RPA-coated ssDNA during recombination-mediated DNA repair to promote nucleation of a RAD51 filament. Our single-molecule imaging of RAD51 filament assembly on its *in vivo* substrate, RPA-coated ssDNA, establishes that a dimer of RAD51 is required for nucleation followed by slow and self-terminating growth. When bound to ssDNA, RPA is a potent and persistent kinetic block to RAD51 binding. The tumor suppressor protein, BRCA2, overcomes this kinetic block by accelerating RAD51 nucleation to rates approaching those seen on ssDNA devoid of RPA. We propose that BRCA2 achieves this task by delivering, via its BRC repeats, a pre-assembled nucleus of RAD51 directly to the DNA. We imagine that the eight BRC repeats of BRCA2 organize RAD51 monomers into a pre-filament that could comprise a nucleus of up to 4 monomers – over-specified, as only two are minimally required – bound to BRC1-4. BRCA2 would also potentially recruit 4 more monomers via BRC5-8 when the BRCA2-RAD51 complex binds to ssDNA (21). The result would be a nascent RAD51 filament of up to 8 monomers – more than one turn of the filament – assembled by the chaperoning capacity of BRCA2. Interestingly, in our experiments containing RPA, RAD51 filaments were restricted to diffraction-limited foci, in contrast to the contiguous structures that form on the ssDNA region in the absence of RPA. This leads us to conclude that, under the conditions of our assay, BRCA2 can facilitate RAD51 nucleation onto RPA-bound ssDNA but it is not capable of stimulating growth of the RAD51 filament beyond spatial limitation of our measurement (∼0.6 kb). *In vivo*, the amount of ssDNA generated following resection or replication fork gap repair may only require a limited stretch (200-300 nt) of RAD51 filament formation to promote strand invasion and homology-directed repair (33). Alternatively, other recombination mediators could assist during the growth phase to further extend RAD51 filaments along regions of ssDNA (15, 16, 34). In *Escherichia coli*, the functional analogs of BRCA2 are RecF, RecO and RecR, which function as a modular set of protein complexes (RecOR and RecFOR), where RecOR stimulates both nucleation and growth of RecA filaments by remodeling the SSB-ssDNA complex, and RecFOR promotes nucleation at or near the ssDNA-dsDNA junctions (28, 35). BRCA2 enhances nucleation of RAD51 filaments, but an as yet-to-be-determined mediator supports RAD51 growth potentially by remodeling RPA-ssDNA structures. Genetically, the RAD51 paralogs are epistatic to BRCA2 (36, 37) and are likely candidates for such a role. Further exploration of this hypothesis awaits more complete biochemical characterization of functionally active human RAD51 paralog proteins.

## Materials and methods

### Single-molecule Measurements

For TIRF imaging, all protein-containing reagents were diluted into single-molecule buffer containing 20 mM TrisOAc (pH 7.5), 50 mM DTT, 20% sucrose, 200 mM NaCl, 2 mM CaCl_2_, 1 mM Mg(OAc)_2_, 1 mM ATP, and 100 nM RPA, unless otherwise indicated. The glass surface was cleaned with piranha solution (7.5% hydrogen peroxide in concentrated sulfuric acid) and functionalized using biotin-PEG silane and blocked. The gapped DNA substrate (gKytos, ∼5 pM molecules) was biotinylated at both ends and attached to the surface *in situ* by alternating the flow off and on using a computer-controlled syringe pump at a flow rate of 4 mL per hour. SYTOX Orange, a dsDNA-specific stain, was used to visualize the dsDNA-regions of the gapped DNA molecules. Owing to the presence of salt and divalent cation, SYTOX Orange dissociated from the dsDNA causing a decrease and eventually disappearance of fluorescence signal. The kinetics of RAD51 nucleation were then observed by injecting 100 nM fluorescent RAD51 (N-terminal linkage with fluorescein, previously described (29)) in the absence or presence of 5 nM 2x-MBP-tagged BRCA2 (purified as previously described (9)).

Epi-fluorescent trapping experiments were performed using a six-channel laminar flow cell. The channels contained the following components, in addition to 20% sucrose and 50 mM DTT: **(Ch 1)** 100 mM NaHCO_3_ (pH 8.3), 100 mM NaCl, 1 mM Mg(OAc)_2_, 5 nM YOYO-1, and 0.2% streptavidin-coated polystyrene beads (1 µm, Bangs); **(Ch 2)** 100 mM NaHCO_3_ (pH 8.3), 100 mM NaCl, 1 mM Mg(OAc)_2_, 50 nM YOYO-1, 2 pM gapped DNA, and 200 nM RPA; **(Ch 3)** 100 mM NaHCO_3_ (pH 8.3), 100 mM NaCl, 1 mM Mg(OAc)_2_, 50 nM YOYO-1, and 50 nM RPA; **(Ch 4)** 100 mM NaHCO_3_ (pH 8.3), and 50 nM RPA; **(Ch 5)** 20 mM Tris(OAc) (pH 7.5), 200 mM NaCl, 2 mM CaCl_2_, 1 mM Mg(OAc)_2_, 1 mM ATP, and 50 nM RPA; and **(Ch 6)** 20 mM Tris(OAc) (pH 7.5), 200 mM NaCl, 2 mM CaCl_2_, 1 mM Mg(OAc)_2_, 1 mM ATP, 50 nM RPA, and 100 nM fluorescent RAD51, unless otherwise indicated. When present, BRCA2 was at 5 nM. Experiments designed to visualize BRCA2 binding, either in the presence or absence of fluorescent RAD51, also contained 1-2 µg/uL α-MBP^AF546^ or 1-2 µg/uL mouse α-BRCA2 (Ab1, Millipore) plus 1-2 µg/µL goat α-mouse IgG^AF546^ (Molecular Probes).

### Total internal reflection fluorescence microscopy and single-channel flow cells

An Eclipse TE2000-U, inverted TIRF microscope (Nikon), using a CFI Plan Apo TIRF 100x, 1.49 N.A., oil-immersion objective was used as previously described(27, 28). Single-channel flow cells were constructed by drilling holes into a glass microscope slide and adhering a coverglass using 3M Thermo-Bond Film (2.5 mil) with a channel cut out from the tape of dimensions (5 mm × 35 mm × 0.1 mm). Inlet ports (PEEK tubing, 0.5 mm inner diameter) were attached to the flow cell using epoxy (Devcon, “5 minute”). Coverglass (Fisher’s *Finest*, 22×50 #1) was cleaned by submersion in Piranha solution (one volume 30% hydrogen peroxide slowly mixed with four volumes of concentrated sulfuric acid) for fifteen to thirty minutes. The coverglass was rinsed with water and then methanol, then sonicated in a solution of 1 M KOH dissolved in methanol for one hour or soaked overnight without sonication. The coverglass was then rinsed with water, methanol, dried under a stream of nitrogen, and then arranged on a slide heater set to 70°C. The surface was functionalized with 330 uL per coverglass (∼0.3 µL per mm^2^) of mPEG-5000-silane (5 mg/mL, Laysan Bio) and biotin-PEG-5000 silane (5 µg/mL, Laysan Bio) dissolved in 0.5 N HCl and 80% ethanol. After the solution evaporated, the top half of a bell-shaped vacuum chamber was placed over the coverglass on the heater and the surface was cured overnight at 70°C under vacuum. The next day, the coverglass was rinsed with water, dried under stream of nitrogen and stored either in a vacuum chamber or in jar purged with nitrogen for up to two weeks. Flow cells were assembled using the double-sided tape with the cut channel. The surface was functionalized by incubating the flow cell successively with the following solutions for 5-10 minutes each: 1) buffer STE (20 mM TrisHCl (pH 7.5), 0.5 mM EDTA and 200 mM NaCl) containing 0.2 mg/mL neutravidin; 2) rinsed with SMB (20 mM TrisOAc (pH 7.5), 50 mM DTT and 20% sucrose); 3) blocked with SMB plus 1.5 mg/mL Roche Blocking Reagent and 2 mg/mL poly-glutamic acid (1500-5500 Da., Sigma-Aldrich); and then 4) rinsed with SMB. The gapped DNA substrate (gKytos, ∼5 pM molecules) was biotinylated at both ends (*see next section*) and attached to the surface *in situ* by alternating the flow off and on using a computer-controlled syringe pump at a flow rate of 4 mL per hour. The dsDNA was visualized by the addition of 200 nM SYTOX Orange (Thermo Fisher).

### Preparation of gapped λ DNA

An engineered derivative of bacteriophage λgt11 containing a λC31 *attP* recognition site was created as previously described(28), and is hereafter called bacteriophage λ*Kytos*. Biotin was incorporated into the *cos* sites of λ*Kytos* in a reaction consisting of 10 mM Tris-HCl (pH 7.9), 10 mM MgCl_2_, 50 mM NaCl, 1 mM DTT, 33 μM dATP, 33 μM dTTP, 33μM dCTP, 33 μM biotin-dGTP, 17 ng/μl λ*Kytos*, and 0.17 U/μl Klenow exo^-^ DNA polymerase. After 15 minutes at 22°C, the polymerase was heat inactivated at 70°C for 20 minutes and the DNA was purified by passing through an S-400 desalting spin column (GE Illustra Microspin). A modified derivative of bacteriophage M13mp7 ssDNA contained the *attB* recognition site, from which a 500 bp dsDNA containing the λC31 *attB* at its center was generated by PCR using Phusion High Fidelity PCR Master Mix from NEB. After heat denaturation, it was annealed to the M13mp7 ssDNA derivative. λC31 integrase was used to recombine λ*Kytos* dsDNA and the annealed 13mp7 ssDNA containing the *attB* recognition site. The construct pHS62, containing the full coding sequence of λC31 integrase, was kindly provided by Margaret Smith(38). Integration of the ssDNA plasmid into λ*Kytos* dsDNA resulted in a gapped λDNA substrate. The gapped λ DNA was purified *in situ* away from non-integrated circular ssDNA either by attaching the molecules to the biotinylated PEG surface for TIRF microscopy, or by attaching the molecules to streptavidin-coated beads during optical trapping experiments.

### Epifluorescence microscopy, optical trapping, and multi-channel flow cells

Optical trapping was achieved on the same TE-2000-U microscope (Nikon) used for the TIRF-based assays. A polarizer (Newport) was used to split the beam from an infrared laser (Spectra-Physics), generating two traps, and a steering mirror (Newport) to control the x-y position of one of the beams(27),(28). Excitation of the sample in epifluorescence mode was achieved using a Cyan 488 nm laser (Picarro) by adjusting the angle of the laser to pass completely through the sample chamber. The fluorescence emission was directed through a dichroic mirror (515/30 nm and 600/40 nm, Chroma). Images were captured on a DU-897E iXon EMCCD camera (Andor). Custom flow cells were constructed as previously described(26, 39). Briefly, the flow cell design (**Fig 1A**) was laser-etched into glass slides (Fisher Scientific 25×75×1 mm) covered with an adhesive abrasive blasting mask (Rayzist Photomask, Inc.) using a 30 Watt Mini-24 Laser Engraver (Epilog Lasers). The slides were sandblasted using 220 grit silicon carbide (Electro Abrasives) to remove residual laser-ablated glass from the channels, resulting in channels ∼100-150 µm deep and 850 µm wide (the total width of the six-channel flow cell was 5.1 mm). Holes were drilled using a diamond-coated bit and a Dremel hand-held drilling tool, washed with 2% Hellmanex III and rinsed with water and methanol. Cover glass (Corning No. 1, 24×60 mm) was cleaned in 1 M KOH/MeOH with sonication for one hour, rinsed with water, methanol, and dried. The cleaned cover glass was attached to the etched microscope slide with UV Optical adhesive #74 (Norland Products) applied through capillary action on a 45°C heat block. The adhesive was cured by placing the flow cell 30 cm from a 100 Watt HBO lamp (Zeiss, Inc.) for 60 minutes followed by curing at 50° C overnight. PEEK tubing with 0.5 mm inner diameter (Upchurch Scientific) was inserted into each of the etched holes to create inlet and outlet connection ports and sealed with epoxy (Devcon, “5 minute”). The flow cell was mounted to the microscope and attached to a computer-controlled syringe pump (KD Scientific). The temperature of the objective lens was held at 37°C by circulating water through a brass and copper collar, machined to fit around the objective lens.

Once DNA dumbbells were assembled, imaging and incubations were performed in the center of the designated channel approximately 100-400 µm downstream of the channel dividers. Experiments designed to visualize BRCA2 binding, either in the presence or absence of fluorescent RAD51, contained 1.5 µg/mL α-MBP^AF546^ (Abcam, labelled with Alexa Fluor 546, Molecular Probes) or 1.5 µg/mL mouse α-BRCA2 (Ab-1, Millipore) plus 1-2 µg/mL goat α-mouse IgG^AF546^ (Molecular Probes) as indicated in the figure legends. Sterile 0.2 µm filtered sucrose solutions were degassed for at least one hour but typically overnight in a vacuum chamber before the addition of 50x buffer and DTT powder, then degassed for an additional 15 minutes; reactions were assembled at room temperature protected from light. Optically trapped molecules were moved between flow channels by movement of the sample stage, which was automated and synchronized with both laser excitation and camera acquisition during dipping experiments using software coded in LabView. Gapped λ DNA molecules containing fluorescent RAD51 filaments were successively transferred from Ch 6 to Ch 5, imaged for 1 s, and immediately transferred back to Ch 6. The time at which a RAD51 cluster first appeared was determined to be the apparent nucleation lag time. Images were processed in ImageJ by frame averaging, background subtraction, and contrast enhancement. Data was analyzed and plotted using GraphPad Prism (v7.0c). Fisher’s exact test was computed using R-Studio (v 0.99.893)

## Supporting information

Movie 1: Binding of BRCA2 on gapped lambda DNA.

Movie 2: RAD51 binding to ssDNA in the absence of RPA using TIRF microscopy.

Movie 3: RAD51 binding to ssDNA in the presence of RPA using TIRF microscopy.

## Acknowledgements

We would like to thank all the members of the Kowalczykowski lab for insightful comments and discussion. J.C.B. was supported in part the NIH-UCD Training Fellowship in Molecular and Cell Biology (T32 GM007377). C.C.D. and J.L.P. were supported by the NIH-UCD T32 Training Program in Oncogenic Signals and Chromosome Biology (CA10052159). J.L.P. was also supported by the NIH F32 Ruth L. Kirschstein National Research Service Award (CA136103). R.B.J. was supported by the American Cancer Society (#IRG 58-012-55); Breast Cancer Alliance; Pilot Project Program grant from Women’s Health Research at Yale, the Yale Comprehensive Cancer Center, and a Liz Tilberis Early Career Award from the Ovarian Cancer Research Fund Alliance. S.C.K. was supported by NIH (GM62653 and GM64745) and DOD-CDMRP (BC171869).

## Supplementary Figure Legends

**Supplementary Figure 1:**
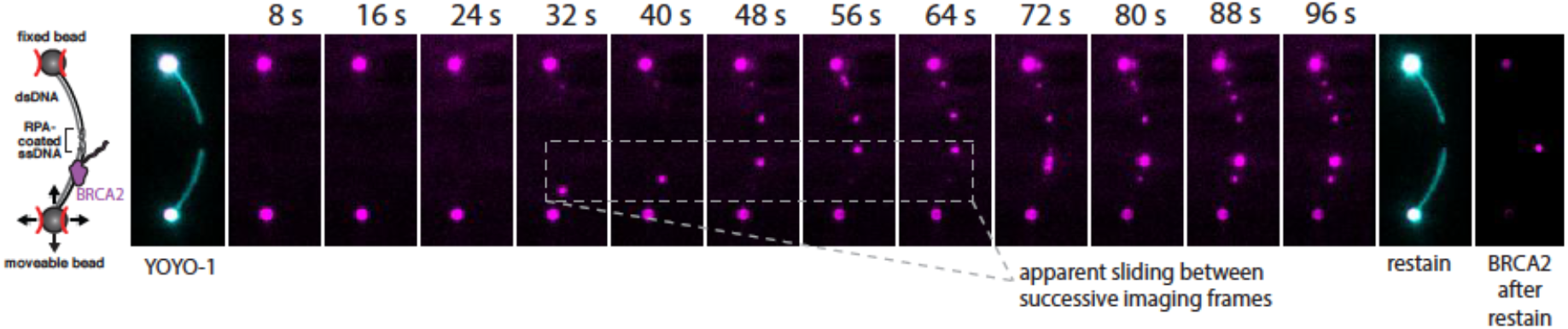
Movement of BRCA2 under flow observed on dsDNA. Montage of a single gapped λDNA molecule held between two optically trapped polystyrene beads in a 6-channel flow cell as described in Figure 1. The molecule was imaged in Channel 5 and iteratively dipped into Channel 6, containing BRCA2 (purple, α-MBP^AF546^), by moving the molecule (*down, then up*) perpendicular to the flow (from *left to right*) between images in the montage. The BRCA2 focus highlighted between 32 s and 64 s slides upwards towards the junction between the dsDNA and RPA-coated ssDNA. At the end of the movie, the gapped λDNA was re-stained in Channel 3, whereupon the less stable BRCA2-dsDNA complexes dissociated.

**Supplementary Figure 2:**
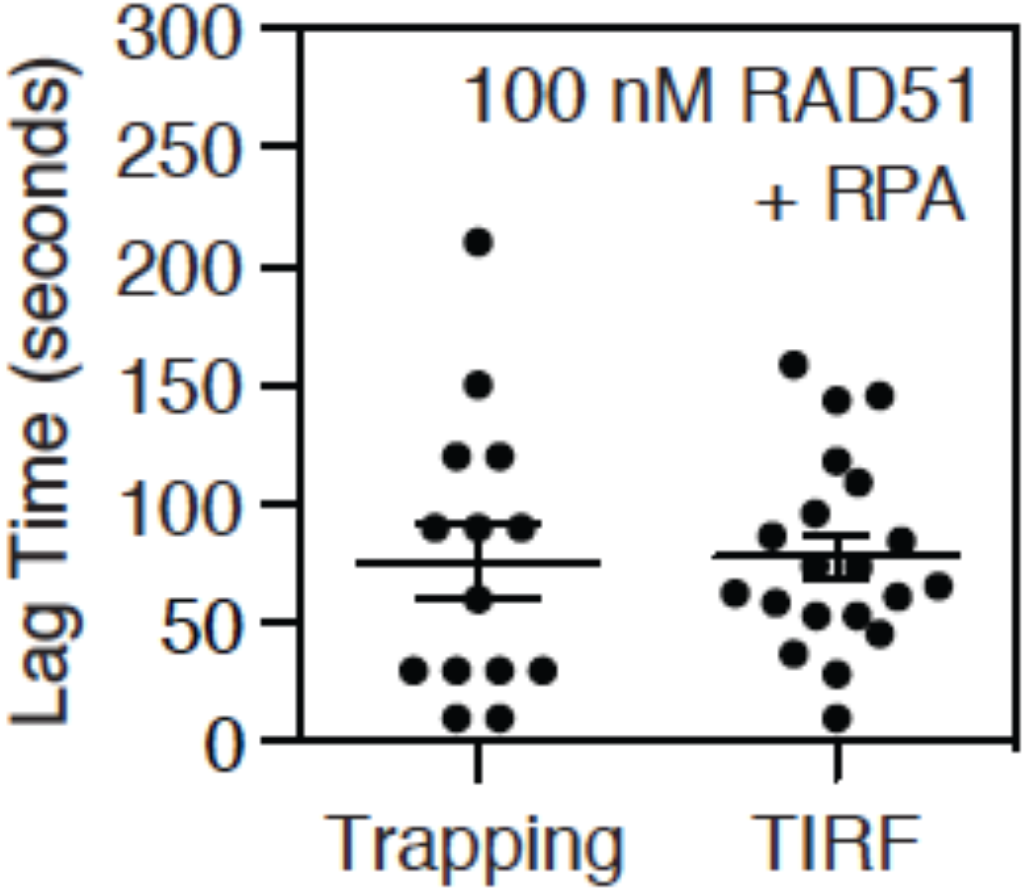
Measurements of RAD51 nucleation using optical trapping and TIRF microscopy are indistinguishable. The lag time was plotted for molecules visualized using either optical trapping as described for Figure 1 (76±16 s, *N=*14) or TIRF as described in Figure 2 (78±9 s, *N=*14) with no observable difference in the distribution of lag times. Error is standard error of the mean.

**Movie 1: Binding of BRCA2 on gapped λDNA**. Movie of a single gapped λDNA molecule held between two optically trapped polystyrene beads in a 6-channel flow cell as described and shown in Supplementary Figure 1. The molecule was imaged in Channel 5 and iteratively dipped into Channel 6, containing BRCA2 (purple), between images in the montage. The BRCA2 appears to slide in the direction if flow (*left to right*) between 32 s and 64 s towards the junction between the dsDNA and RPA-coated ssDNA. At the end of the movie, the gapped λDNA was re-stained in Channel 3.

**Movie 2: RAD51 binding to ssDNA in the absence of RPA using TIRF microscopy**. A single gapped λDNA molecule was initially visualized using SYTOX Orange (*red*), which was subsequently dissociated upon the addition of binding buffer and fluorescent RAD51 (*green*) in the absence of RPA. Movie was collected as described in **Fig. 2A** and corresponds to the montage shown in **Fig. 2B**.

**Movie 3: RAD51 binding to ssDNA in the presence of RPA using TIRF microscopy**. A single gapped λDNA molecule was initially visualized using SYTOX Orange (*red*), which was subsequently dissociated upon the addition of binding buffer and fluorescent RAD51 (*green*) in the presence of RPA. Movie was collected as described in **Fig. 2A** and corresponds to the montage shown in **Fig. 2C**.

## References

1. R. Wooster et al., Identification of the breast cancer susceptibility gene BRCA2. Nature 378, 789–792 (1995).

2. R. Wooster et al., Localization of a breast cancer susceptibility gene, BRCA2, to chromosome 13q12-13. Science 265, 2088–2090 (1994).

3. A. K. Wong, R. Pero, P. A. Ormonde, S. V. Tavtigian, P. L. Bartel, RAD51 interacts with the evolutionarily conserved BRC motifs in the human breast cancer susceptibility gene brca2. J. Biol. Chem. 272, 31941–31944 (1997).

4. S. K. Sharan et al., Embryonic lethality and radiation hypersensitivity mediated by Rad51 in mice lacking Brca2. Nature 386, 804–810 (1997).

5. V. P. Yu et al., Gross chromosomal rearrangements and genetic exchange between nonhomologous chromosomes following BRCA2 inactivation. Genes Dev. 14, 1400–1406 (2000).

6. M. E. Moynahan, A. J. Pierce, M. Jasin, BRCA2 is required for homology-directed repair of chromosomal breaks. Mol. Cell 7, 263–272 (2001).

7. S. S. Yuan et al., BRCA2 is required for ionizing radiation-induced assembly of Rad51 complex in vivo. Cancer Res. 59, 3547–3551 (1999).

8. B. C. Godthelp, F. Artwert, H. Joenje, M. Z. Zdzienicka, Impaired DNA damage-induced nuclear Rad51 foci formation uniquely characterizes Fanconi anemia group D1. Oncogene 21, 5002–5005 (2002).

9. R. B. Jensen, A. Carreira, S. C. Kowalczykowski, Purified human BRCA2 stimulates RAD51-mediated recombination. Nature 467, 678–683 (2010).

10. P. L. Welcsh, K. N. Owens, M. C. King, Insights into the functions of BRCA1 and BRCA2. Trends Genet. 16, 69–74 (2000).

11. J. Liu, T. Doty, B. Gibson, W. D. Heyer, Human BRCA2 protein promotes RAD51 filament formation on RPA-covered single-stranded DNA. Nat Struct Mol Biol 17, 1260–1262 (2010).

12. T. Thorslund et al., The breast cancer tumor suppressor BRCA2 promotes the specific targeting of RAD51 to single-stranded DNA. Nat Struct Mol Biol 17, 1263–1265 (2010).

13. R. Prakash, Y. Zhang, W. Feng, M. Jasin, Homologous recombination and human health: the roles of BRCA1, BRCA2, and associated proteins. Cold Spring Harb Perspect Biol 7, a016600 (2015).

14. A. Zelensky, R. Kanaar, C. Wyman, Mediators of homologous DNA pairing. Cold Spring Harb Perspect Biol 6, a016451 (2014).

15. J. C. Bell, S. C. Kowalczykowski, Mechanics and Single-Molecule Interrogation of DNA Recombination. Annu. Rev. Biochem. 85, 193–226 (2016).

16. S. C. Kowalczykowski, An Overview of the Molecular Mechanisms of Recombinational DNA Repair. Cold Spring Harb Perspect Biol 7, a016410 (2015).

17. P. Bork, N. Blomberg, M. Nilges, Internal repeats in the BRCA2 protein sequence. Nat. Genet. 13, 22–23 (1996).

18. G. Bignell, G. Micklem, M. R. Stratton, A. Ashworth, R. Wooster, The BRC repeats are conserved in mammalian BRCA2 proteins. Hum. Mol. Genet. 6, 53–58 (1997).

19. L. Pellegrini et al., Insights into DNA recombination from the structure of a RAD51-BRCA2 complex. Nature 420, 287–293 (2002).

20. A. Carreira et al., The BRC repeats of BRCA2 modulate the DNA-binding selectivity of RAD51. Cell 136, 1032–1043 (2009).

21. A. Carreira, S. C. Kowalczykowski, Two classes of BRC repeats in BRCA2 promote RAD51 nucleoprotein filament function by distinct mechanisms. Proc. Natl. Acad. Sci. U. S. A. 108, 10448–10453 (2011).

22. H. Yang et al., BRCA2 function in DNA binding and recombination from a BRCA2-DSS1-ssDNA structure. Science 297, 1837–1848 (2002).

23. S. C. Kowalczykowski, Molecular mimicry connects BRCA2 to Rad51 and recombinational DNA repair. Nat. Struct. Biol. 9, 897–899 (2002).

24. W. Zhao et al., Promotion of BRCA2-Dependent Homologous Recombination by DSS1 via RPA Targeting and DNA Mimicry. Mol. Cell 59, 176–187 (2015).

25. G. Chatterjee, J. Jimenez-Sainz, T. Presti, T. Nguyen, R. B. Jensen, Distinct binding of BRCA2 BRC repeats to RAD51 generates differential DNA damage sensitivity. Nucleic Acids Res. 44, 5256–5270 (2016).

26. A. L. Forget, C. C. Dombrowski, I. Amitani, S. C. Kowalczykowski, Exploring protein-DNA interactions in 3D using in situ construction, manipulation and visualization of individual DNA dumbbells with optical traps, microfluidics and fluorescence microscopy. Nat Protoc 8, 525–538 (2013).

27. A. L. Forget, S. C. Kowalczykowski, Single-molecule imaging of DNA pairing by RecA reveals a three-dimensional homology search. Nature 482, 423–427 (2012).

28. J. C. Bell, J. L. Plank, C. C. Dombrowski, S. C. Kowalczykowski, Direct imaging of RecA nucleation and growth on single molecules of SSB-coated ssDNA. Nature 491, 274–278 (2012).

29. J. Hilario, I. Amitani, R. J. Baskin, S. C. Kowalczykowski, Direct imaging of human Rad51 nucleoprotein dynamics on individual DNA molecules. Proc. Natl. Acad. Sci. U. S. A. 106, 361–368 (2009).

30. H. Naora, H. Naora, M. Izawa, V. G. Allfrey, A. E. Mirsky, Some observations on differences in composition between the nucleus and cytoplasm of the frog oocyte. Proc. Natl. Acad. Sci. U. S. A. 48, 853–859 (1962).

31. R. Galletto, I. Amitani, R. J. Baskin, S. C. Kowalczykowski, Direct observation of individual RecA filaments assembling on single DNA molecules. Nature 443, 875–878 (2006).

32. A. Candelli et al., Visualization and quantification of nascent RAD51 filament formation at single-monomer resolution. Proc. Natl. Acad. Sci. U. S. A. 111, 15090–15095 (2014).

33. L. S. Symington, End resection at double-strand breaks: mechanism and regulation. Cold Spring Harb Perspect Biol 6, a016436 (2014).

34. J. C. Bell, S. C. Kowalczykowski, RecA: Regulation and Mechanism of a Molecular Search Engine. Trends Biochem. Sci. 41, 491–507 (2016).

35. K. Morimatsu, S. C. Kowalczykowski, RecFOR proteins load RecA protein onto gapped DNA to accelerate DNA strand exchange: a universal step of recombinational repair. Mol. Cell 11, 1337–1347 (2003).

36. R. B. Jensen, A. Ozes, T. Kim, A. Estep, S. C. Kowalczykowski, BRCA2 is epistatic to the RAD51 paralogs in response to DNA damage. DNA Repair (Amst) 12, 306–311 (2013).

37. J. Chun, E. S. Buechelmaier, S. N. Powell, Rad51 paralog complexes BCDX2 and CX3 act at different stages in the BRCA1-BRCA2-dependent homologous recombination pathway. Mol. Cell. Biol. 33, 387–395 (2013).

38. H. M. Thorpe, M. C. Smith, In vitro site-specific integration of bacteriophage DNA catalyzed by a recombinase of the resolvase/invertase family. Proc. Natl. Acad. Sci. U. S. A. 95, 5505–5510 (1998).

39. I. Amitani, B. Liu, C. C. Dombrowski, R. J. Baskin, S. C. Kowalczykowski, Watching individual proteins acting on single molecules of DNA. Methods Enzymol. 472, 261–291 (2010).

